# Second-order kinetics describe systemic clearance of therapeutic bacteriophages

**DOI:** 10.64898/2026.08.05.742954

**Authors:** Joël L. Gerber, David R. Cameron, Josef Prazak, Bülent Gözel, Camille Boross, Guido Stirnimann, Grégory Resch, Yok-Ai Que

## Abstract

Bacteriophage therapy is a promising alternative to antibiotics, yet its clinical translation is limited by the lack of a quantitative pharmacological framework to guide dosing and to predict efficacy. Here, we define the pharmacokinetics of therapeutic phages using a rat tissue cage model, which allows parallel sampling from blood and an artificial interstitial compartment. Across five virulent phages of three morphotypes targeting two pathogens, systemic clearance consistently followed second-order, concentration-dependent kinetics, representing a paradigmatic shift from frequently assumed first-order models. Phages rapidly distributed to peripheral compartments, where exposure was strongly influenced by administration route. Intravenous delivery maximized systemic titers but limited peripheral exposure, whereas local administration achieved high concentrations at target sites with undetectable systemic redistribution. Repeated dosing enhanced exposure but not peak titers. These findings define fundamental parameters to establish a quantitative framework for phage pharmacokinetics and support rational dose design.

## INTRODUCTION

The accelerating rise of multidrug-resistant bacteria^1,2^ is outpacing the development of new antibiotics^3,4^, creating an urgent need for alternative anti-infective strategies. Bacteriophage therapy, the use of viruses that specifically infect and lyse bacteria, has re-emerged as a promising approach^5,6^. Despite strong bactericidal activity observed in vitro^6,7^, encouraging results in animal models^8^, and growing clinical interest^9,10^, efficacy has yet to be consistently demonstrated in randomized controlled trials^11–13^.

A central limitation is the absence of a quantitative pharmacological framework to guide dosing and treatment schedules^14,15^. Unlike antibiotics, for which development has been shaped by rigorous pharmacokinetic/pharmacodynamic (PK/PD) studies^16,17^, phage therapy has largely adopted dosing strategies supported by limited PK data. As a result, phage administration remains highly heterogeneous and mostly empirical, limiting comparability across studies and potentially contributing to inconclusive clinical trial outcomes. Notably, systemic phage kinetics are frequently interpreted and modelled using first-order kinetics^18–22^, implicitly assuming constant half-life by analogy to small-molecule drugs, despite limited experimental validation^23^. In practice however, circulating phage titers often exhibit rapid, exponential decay that deviates from these assumptions^24^, suggesting that classical drug-based PK models may not adequately describe phage behavior in vivo.

This discrepancy reflects a more fundamental issue; phages are not conventional drugs. Their in vivo behavior is shaped by dynamic interactions with host biology, including uptake by the reticuloendothelial system^25,26^, immune recognition^27^, and potential amplification at sites of infection^28^. These processes introduce complex behaviors that may not adequately be captured by classical PK models. Consequently, key PK parameters, including clearance are incompletely defined, limiting the rational design of dosing strategies and treatment schedules. Moreover, the relationship between intravenous administration and phage exposure at the site of infection remains insufficiently understood, possibly contributing to the predominant use of local or topical administration routes in phage therapy^29,30^.

Here, we sought to establish a mechanistic framework for in vivo phage PK. Experimental models specifically designed for the study of bacteriophage absorption, distribution and elimination remain scarce^23,31^. Consequently, most *in vivo* studies have relied on murine infection models adapted from antibiotic efficacy testing and have primarily focused on therapeutic outcomes rather than phage PK^23^. These models were not designed to enable serial sampling at peripheral sites of infection, as they were originally developed to investigate systemic PK/PD relationships for antibiotics. In addition, the limited circulating blood volume of mice precludes repeated blood sampling, restricting longitudinal assessment of phage clearance from circulation.

Using an adapted rat tissue cage model, we quantified systemic clearance, distribution to infection sites, and exposure dynamics of diverse phages with therapeutic potential targeting the priority pathogens *Staphylococcus aureus* and *Pseudomonas aeruginosa*. By integrating experimental data with kinetic modeling, we demonstrate that systemic phage elimination follows second-order kinetics, revealing a concentration-dependent clearance mechanism conserved across phages. We further show that administration route and dosing frequency differentially shape systemic and peripheral exposure profiles. Together, these findings provide a quantitative framework for phage pharmacology and establish principles for rational dose design to optimize targeting of bacterial pathogens.

## RESULTS

### Rat tissue cage model enables comprehensive phage pharmacokinetic analyses

We developed a rat model dedicated to the study of phage pharmacology (Figure 1). A totally implantable venous access port (TIVAP) was inserted to allow serial venous blood sampling (central or systemic compartment) and repeated intravenous phage administration Additionally, two tissue cages (TCs) were implanted subcutaneously to create artificial peripheral compartments that could be repeatedly sampled for pharmacokinetic analyses (Figure 1A). After surgery, animals recovered for three weeks to allow implant encapsulation and tissue integration, resulting in compartmentalization and accumulation of interstitial fluid, i.e. formation of an artificial seroma, within each TC (Figure 1B). A conceptual pharmacokinetic model is illustrated in Figure 1C, with the bloodstream representing the central compartment and the TCs representing peripheral compartments corresponding to potential infection sites.

**FIGURE 1:**
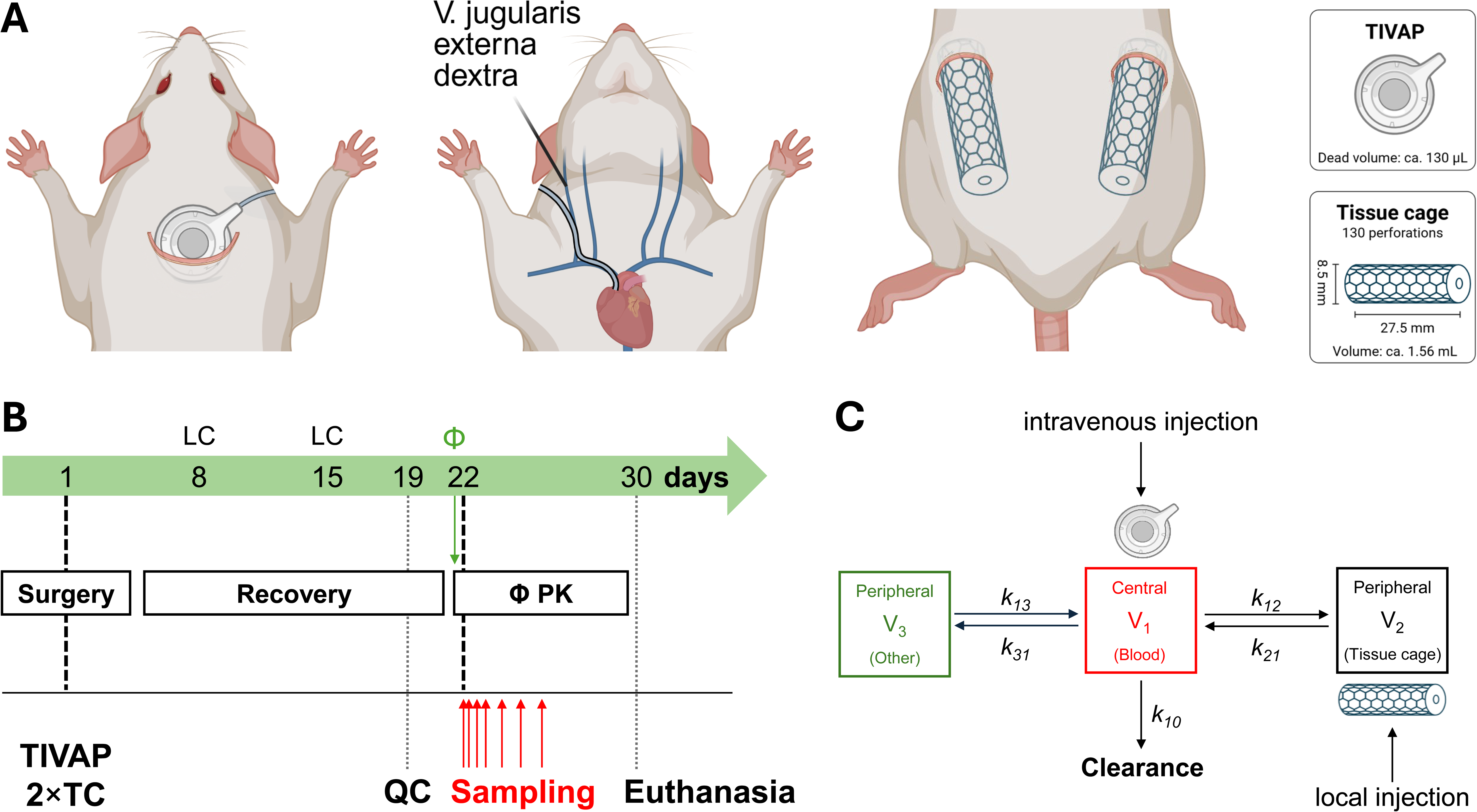
Experimental platform for bacteriophage pharmacokinetics in rats. (**A**) Schematic of the surgical setup. Rats were equipped with a total implantable venous access port (TIVAP) via the right jugular vein to enable repeated intravenous administration and serial blood sampling. In addition, two cylindric, perforated perfluoroalkoxy alkane tissue cages (TC) were implanted subcutaneously on the dorsum to create an artificial, puncturable peripheral compartment. (**B**) Study timeline. After surgery (day 1), animals underwent a recovery phase, during which lock was changed twice (LC, day 8 and day 15), before phage pharmacokinetic (Φ PK) experiments. Sampling points included sterility/quality control (QC, day 19), serial blood and tissue cage fluid sampling, before euthanasia. (**C**) Conceptual pharmacokinetic model. Phages were administered intravenously via the TIVAP (central compartment, V_1_, blood), locally into the tissue cage (peripheral compartment, V_2_), intraperitoneally or subcutaneously (both V_3_). Exchange between compartments is described by rate constants (k_12_, k_21_), with an additional peripheral compartment (V_3_) representing the other compartments, including fat tissue and the peritoneal cavity (k_13_, k_31_). Systemic clearance is represented by k_10_.

### Systemic phage clearance follows second-order kinetics

To characterize systemic phage elimination, rats received a single intravenous dose (5 × 10^9^ PFU) of one of five virulent bacteriophages with distinct morphologies targeting either *Staphylococcus aureus* (ΦK, Φ2002, and Φ44AHJD) or *Pseudomonas aeruginosa* (ΦPB1 and ΦDMS3*vir*). Following administration, circulating phage titers peaked within five minutes and then rapidly declined for all phages (Figure 2). Peak concentrations varied between phages, reaching 1.85 × 10^7^ PFU/mL on average (n = 32), followed by a decay that did not linearize after logarithmic transformation, as would have been expected for first-order elimination kinetics (Supplemental Figure 1). By 48h, titers had decreased to a mean trough concentration of 1.47 × 10^8^ PFU/mL. Importantly, this kinetic profile was conserved across phages with distinct morphology, taxonomy, and bacterial host specificity, suggesting a common host-driven clearance mechanism.

**FIGURE 2:**
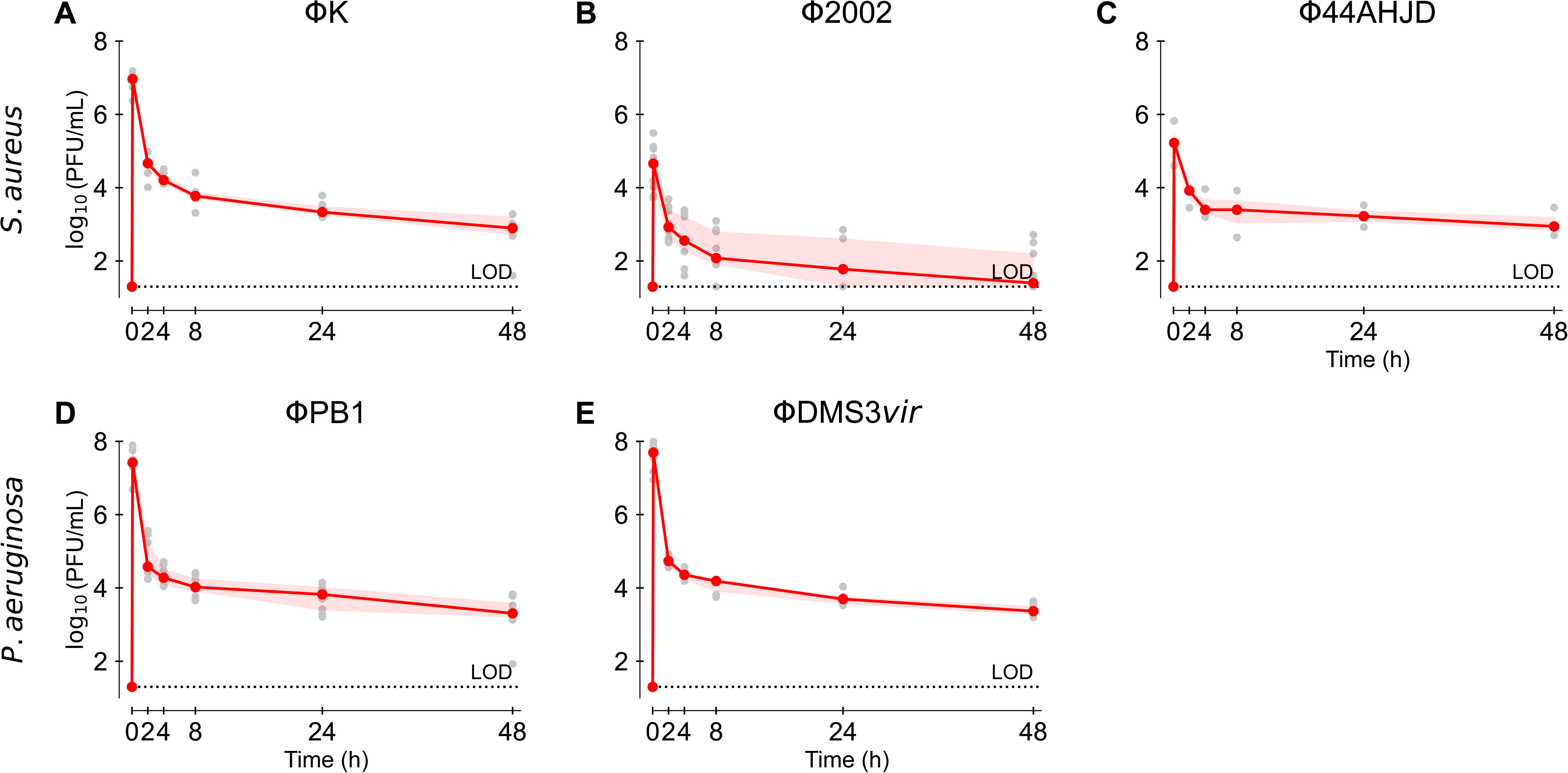
Systemic pharmacokinetics of bacteriophages following a single intravenous dose. Blood phage titers were quantified over 48 h after a single intravenous administration (5 × 10^9^ plaque forming units [PFU]) via the TIVAP. Kinetic profiles are shown for *Staphylococcus aureus* phages ΦK (**A**; n = 6), Φ2002 (**B**; n = 9), and Φ44AHJD (**C**; n = 3), and *Pseudomonas aeruginosa* phages ΦPB1 (**D**; n = 8) and ΦDMS3*vir* (**E**; n = 6). Individual animals are represented by gray points; red lines indicate the median, and shaded areas denote the interquartile range (IQR). The dotted horizontal line indicates the limit of detection (LOD; 20 PFU/mL).

To determine the underlying elimination kinetics, observed data were compared to zero-order (Figure 3, first column), first-order (Figure 3, second column), and second-order models (Figure 3, third column). The theoretical properties of each kinetic order, including rate laws, half-life behavior, and linearization strategies, are summarized in Supplemental Figure 1 and Supplemental Table 1. Empirically, phage decay consistently failed to conform to archetypal elimination models that describe small molecule PK, including first-order kinetics (constant half-life, Supplemental Figure 1BEH), and zero-order kinetics (concentration-independent elimination at a constant rate, Supplemental Figure 1ADG). Instead, the data closely followed second-order kinetics, characterized by a concentration-dependent clearance rate (Equation 1).

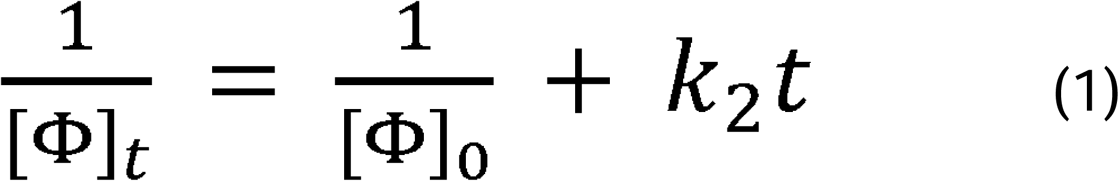

**FIGURE 3:**
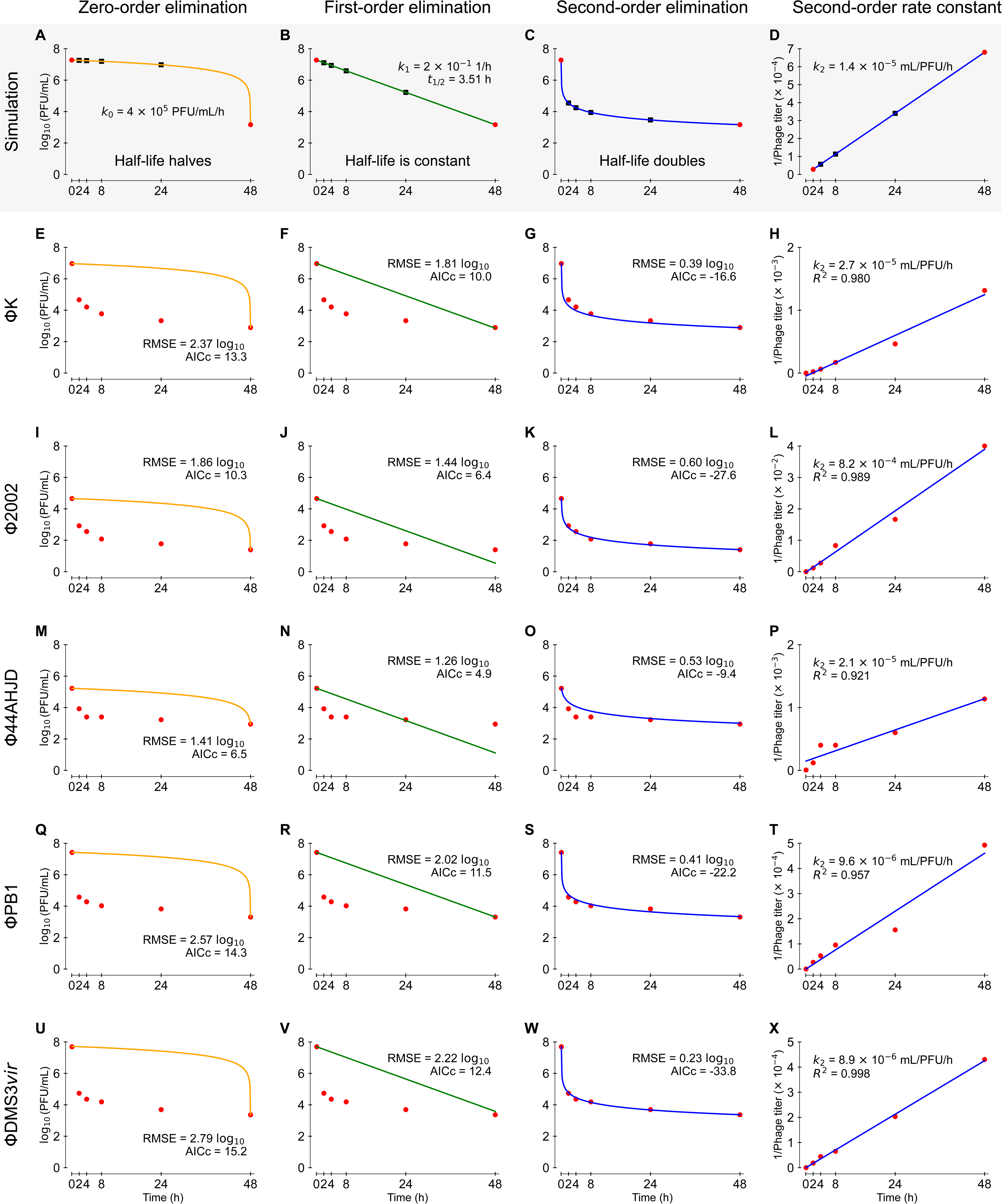
Systemic phage clearance is best described by second-order elimination kinetics. Simulated elimination profiles under different kinetic order assumptions. Phage decay was simulated over 48 h starting from 1.85 × 10^7^ plaque forming unit (PFU)/mL (mean peak across all profiles, n = 32) and ending to 1.47 × 10³ PFU/mL (mean trough). Black data points represent simulated values at defined time points. Zero-order elimination (**A**, orange) assumes a constant rate of decline with halving of the half-life; first-order elimination (**B**, green) assumes a constant half-life; and second-order elimination (**C**, blue) assumes half-life doubling. Panel (**D**) shows the linear relationship between inverse phage titer and time in second-order kinetics, where the slope of the linear regression corresponds to the second-order rate constant (*k_2_*). (**E–I**) Model fits (colored lines) to observed median titers (red datapoints) for individual phages after a single intravenous administration of 5 × 10^9^ PFU: ΦK **(E**), Φ2002 (**F**), Φ44AHJD (**G**), ΦPB1 (**H**), and ΦDMS3*vir* (**I**). For each phage, zero-order (orange), first-order (green), and second-order (blue) fits are shown. The corresponding linear regressions of inverse phage titer versus time are displayed in blue, with *k_2_* specifying the second-order rate constant (slope) and *R^2^* showing regression fit. The root mean square error (RMSE) and corrected Akaike information criterion (AICc), calculated from log_10_-transformed titers, indicate the average prediction error and relative model fit, respectively.

Transformation of the phage titer data to inverse concentration (1/[Φ]) over time revealed strong linear relationships with *R^2^* values ranging from 0.921 to 0.998 across all phages (Figure 3), consistent with second-order elimination kinetics. The derived second-order rate constants (*k_2_*) varied between phages, indicating phage-specific elimination dynamics analogous to compound-specific half-lives in first-order kinetics. Comparison of predefined kinetic models using root mean square error demonstrated superior performance of the second-order model over zero-and first-order models. Similarly, analysis of phage kinetic data extracted from four independent studies^18,20,32,33^ (Supplemental Table 2) were better described by a second-order elimination model than by conventional first-order kinetics, irrespective of mammalian species, sex or phage, with remarkably similar rate constants (Figure 4, Supplemental Figure 2, Supplemental Table 2). Together, these findings indicate that systemic phage clearance follows second-order kinetics, with elimination accelerating at high titers and slowing as concentrations decrease. Accordingly, the apparent half-life is not constant but inversely related to phage concentration, as predicted for second-order kinetics (Equation 2).

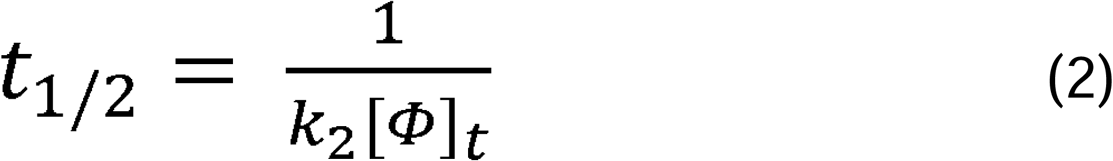

**FIGURE 4:**
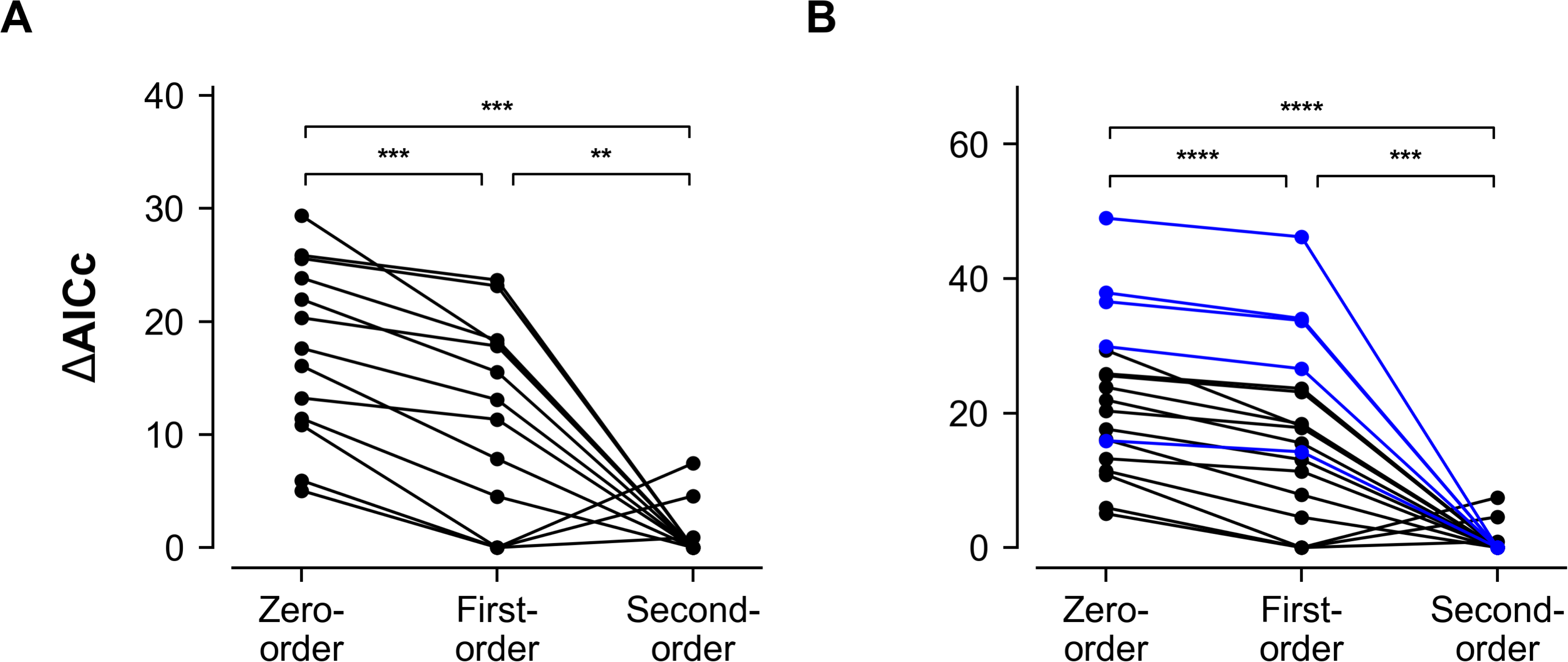
Comparison of model fit by differences in the corrected Akaike information criterion. Differences in the corrected Akaike information criterion (ΔAICc) values among zero-, first-, and second-order elimination models are shown for (**A**) published datasets (black; n = 13) or (**B**) the combined published and generated (blue; n = 5) datasets (total n = 18), where ΔAICc = 0 indicates the best-fitting model. Overall ΔAICc differed significantly between models (*p* < 0.001 each). Pairwise comparisons are indicated by asterisks: *p* < 0.05 (*), *p* < 0.01 (**), *p* < 0.001 (***), and *p* < 0.0001 (****).

### Phages distribute rapidly to peripheral compartments after intravenous administration

Following intravenous administration, phages rapidly distributed from the systemic circulation into the tissue cage fluid (TCF), used here as a surrogate for peripheral infection sites (Figure 5). Detectable titers in TCF appeared within five minutes and reached peak levels approximately one to two orders of magnitude lower than in blood, before approaching equilibrium between both compartments from 2h onward. Overall, the TCF concentration–time profiles closely paralleled systemic kinetics, consistent with rapid exchange between central and peripheral compartments as described in the conceptual model (Figure 1C).

**FIGURE 5:**
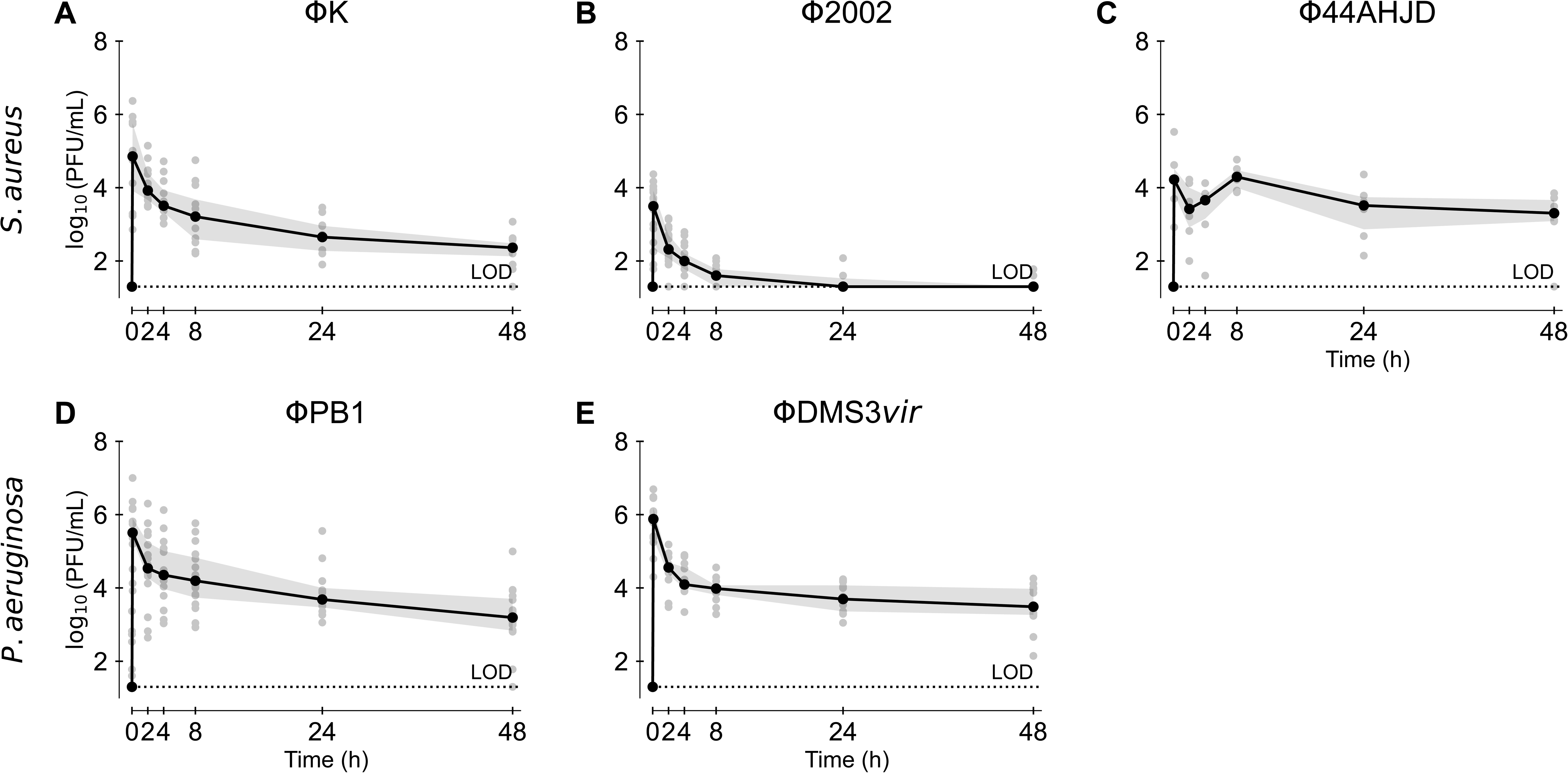
Local phage kinetics in tissue cage fluid following a single intravenous dose. Phage titers in tissue cage fluid were measured over time after a single intravenous administration of 5 × 10^9^ plaque forming units (PFU). Kinetic profiles are shown for ΦK (**A**; n = 12), Φ2002 (**B**; n = 18), Φ44AHJD (**C**; n = 6), ΦPB1 (**D**; n = 16), and ΦDMS3*vir* (**E**; n = 12). Individual measurements are shown as gray points. Black lines represent median titers, and shaded areas indicate the interquartile range (IQR). The dotted horizontal line indicates the limit of detection (LOD; 20 PFU/mL).

### Injection route determines systemic and peripheral phage exposure

Subsequent absorption and distribution studies focused on ΦK. To assess the impact of administration route, ΦK was delivered intravenously, intraperitoneally or subcutaneously (5 × 10^9^ PFU), or locally directly into the TC at a lower concentration allowing for direct comparison (5 × 10^7^ PFU, Figure 6; Supplemental Figure 3). Marked differences in both systemic and peripheral exposure were observed across routes.

**FIGURE 6:**
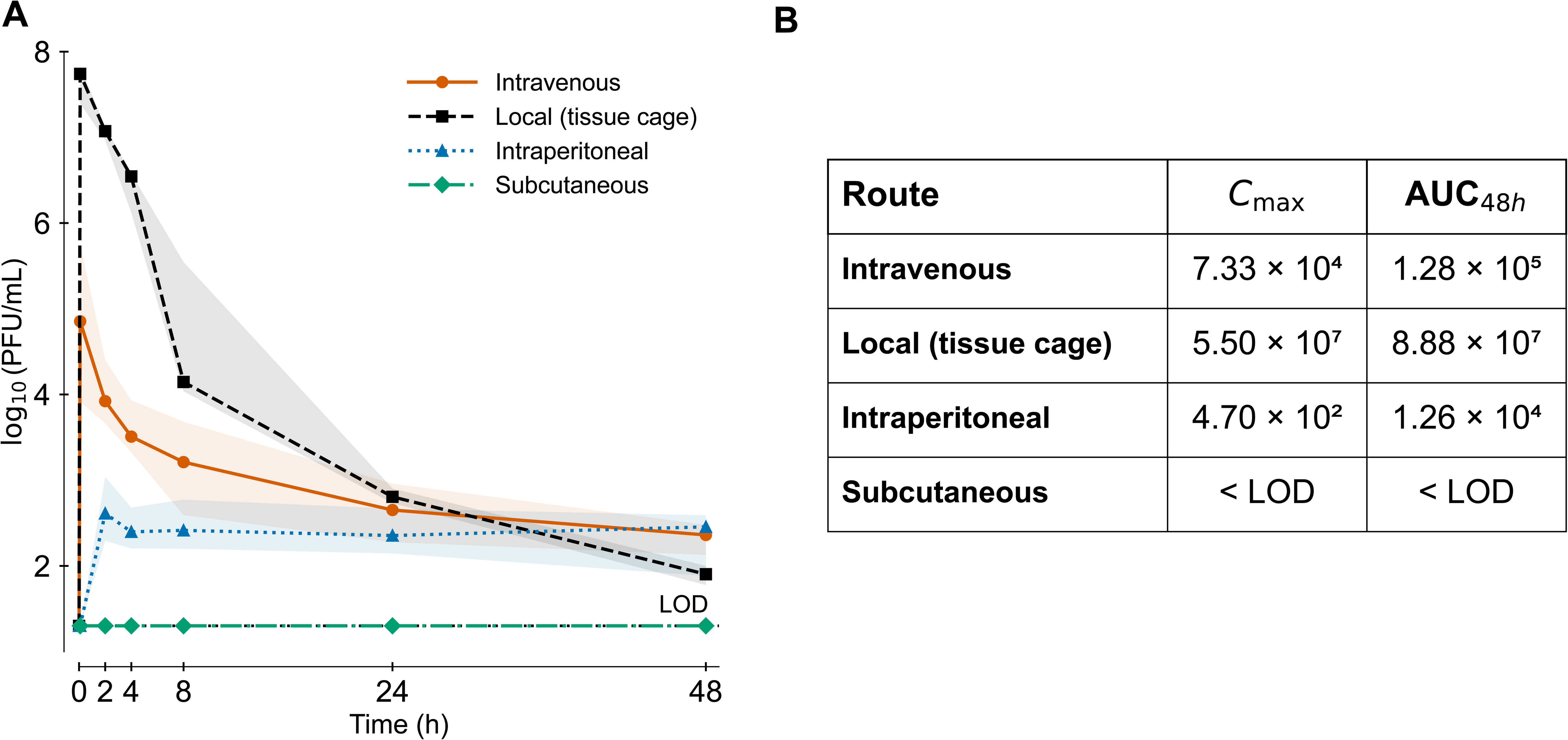
Route-dependent phage exposure in tissue cage fluid following single-dose administration of ΦK. (**A**) Phage titers in tissue cage fluid (TCF) following administration of ΦK via different routes: intravenous (n = 12), local (tissue cage; n = 3), intraperitoneal (n = 4), and subcutaneous (n = 2). Doses were 5 × 10^9^ plaque forming units (PFU) for intravenous administration and 5 × 10^7^ PFU for local tissue cage administration. Data points represent median titers, and shaded areas indicate the interquartile range (IQR). The dotted horizontal line indicates the limit of detection (LOD; 20 PFU/mL). (**B**) Maximum concentration (C_max_), and systemic exposure (AUC_48h_) in TCF calculated from median titers over 48 h using the trapezoidal rule.

Intravenous administration produced the highest systemic titers, with rapid peak blood concentrations (C_max_ 9.25 × 10^6^ PFU/ml), and the greatest overall systemic exposure over 48 hours (expressed as area under the curve, AUC_48h_ 9.50 × 10^6^ PFU×h/ml; Supplemental Figure 3). In contrast, intraperitoneal administration resulted in substantially lower systemic exposure (C_max_ ∼10^4^ PFU/mL, AUC_48h_ 1.31 × 10^5^ PFU×h/ml), consistent with limited bioavailability, whereas subcutaneous administration produced no detectable systemic titers.

Conversely, local intra-TC administration yielded the highest phage concentrations within the peripheral compartment (C_max_ 5.50 × 10^7^ PFU/mL, AUC_48h_ 8.88 × 10^7^ PFU×h/ml; Figure 6). Intravenous administration resulted in substantially lower peripheral exposure (C_max_ 7.33 × 10^4^ PFU/mL, AUC_48h_ 1.28 × 10^5^ PFU×h/ml), whereas intraperitoneal and subcutaneous injections did not produce relevant peripheral titers.

### Repeated dosing increases exposure but not peak titers

Although phage therapy is commonly administered as repeated doses or through combined routes, experimental evidence supporting these strategies remains limited. To evaluate cumulative effects, ΦK was administered at 0, 4, 8, and 24h either intravenously (5 × 10^9^ PFU), locally by intra-TC injection (5 × 10^7^ PFU) or through combined administration routes (Figure 7). Despite repeated intravenous dosing, systemic peak concentrations remained comparable across injections, indicating no accumulation in the central compartment (Figure 7A). This behavior is consistent with second-order kinetics, in which clearance is concentration-dependent and accelerates at higher concentrations, in contrast to first-order elimination, as illustrated by modeling (Figure 7D and 7E). Notably, first-order simulations predicted systemic accumulation, with a 1.7-fold increase in peak titers after repeated dosing (Figure 7D), although this effect appeared attenuated on a logarithmic scale. Local intra-TC administration, alone or combined with intravenous dosing, produced substantially higher peripheral phage titers, largely driven by the local route, with similar peak and trough concentrations over time, suggesting minimal peripheral accumulation (Figure 7B and 7C).

**FIGURE 7:**
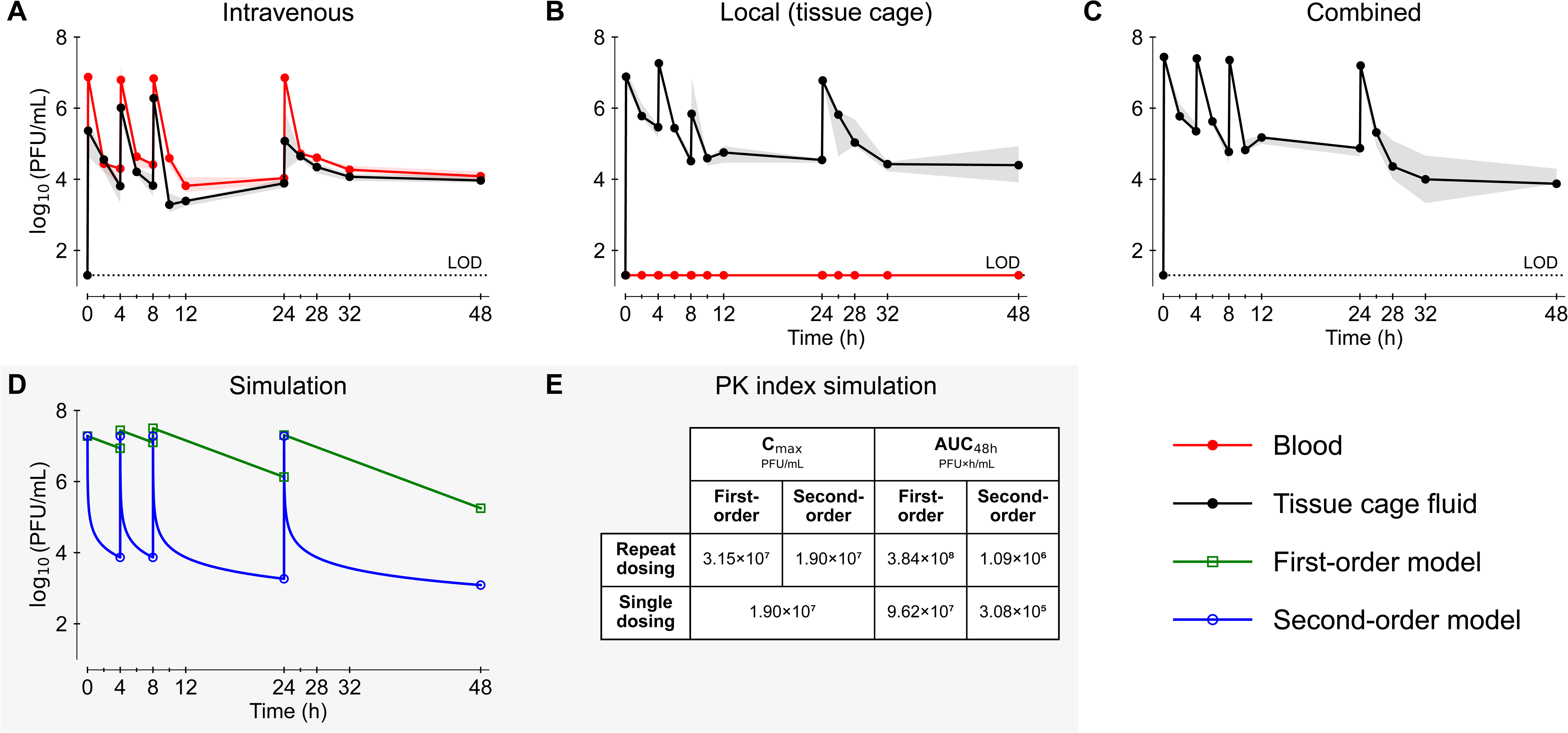
Repeat dosing of ΦK reveals accumulation consistent with second-order kinetics. (**A–C**) Blood (red) and tissue cage fluid titers (black) following repeated administration of ΦK via intravenous (**A**), local (tissue cage; **B**), or combined intravenous and local routes (**C**) (n = 3 per group). Doses were 5 × 10^9^ plaque forming units (PFU) per injection for intravenous administration and 5 × 10^7^ PFU per injection for local administration, given at 0, 4, 8, and 24 h. Red points indicate blood titers and black points indicate TCF titers; lines represent medians, and shaded areas indicate the interquartile range (IQR). The dotted horizontal line indicates the limit of detection (LOD; 20 PFU/mL). (**D**) Simulated concentration-time profiles comparing first-order (green) and second-order (blue) elimination kinetics using parameters derived from Figure 3 (mean peak titers, half-life, and *k_2_*). (**E**) Simulated pharmacokinetic indices, including maximum concentration (C_max_) and exposure (AUC_48h_), for single versus repeat dosing under first-and second-order models.

Simulations based on parameters derived from second-order modeling (Figure 7E) further supported these observations, showing that repeated dosing under second-order kinetics leads to improved peripheral exposure without significant systemic accumulation. Although systemic C_max_ remained stable, cumulative peripheral exposure metrics, including AUC_48h_ and time above threshold concentration, increased substantially.

## DISCUSSION

Advancing phage therapy requires a conceptual framework linking PK to rational treatment design. Using a dedicated rat tissue cage model, we defined the *in vivo* PK of therapeutic phages in systemic and peripheral compartments and identified key determinants governing elimination, distribution, and accumulation for quantitative modeling. Across five structurally and taxonomically distinct virulent phages targeting *S. aureus* and *P. aeruginosa*, systemic pharmacokinetic behavior was highly conserved and consistently followed second-order elimination kinetics, independent of phage family, morphology, or host specificity.

Zero-and first-order elimination kinetics underpin classical small molecule pharmacokinetics, with non-linear Michaelis-Menten kinetics describing the saturation-dependent transition between the two^34,35^. In phage PK, first-order elimination has likewise been widely assumed by extrapolation from the small-molecule paradigm^18–22^. However, the elimination profiles we observed did not conform to either of the two classical models. Given that phage clearance from the bloodstream primarily involves the reticuloendothelial system^25,26^, rather than the renal and biliary excretion pathways that drive small-molecule elimination, it is not surprising that conventional PK models inadequately capture phage behavior in vivo. Instead, our rat data demonstrate that second-order kinetics, a well-established principle in chemical (reaction) kinetics but not small-molecule PK, more accurately captures both the magnitude and trajectory of systemic phage decay and predicts circulating infectious phage concentrations. Importantly, re-analysis of published phage PK datasets (n = 13) converge on the same conclusion, indicating that systemic phage elimination follows second-order elimination across diverse phages, mammalian species, and experimental settings. Together, these findings identify second-order kinetics as a more appropriate quantitative framework for future PK/PD modelling and dose optimization.

Predictive PK/PD indices of phage therapy efficacy, analogous to those used to guide antibiotic treatment (time>MIC, AUC/MIC, or C_max_/MIC), are yet to be established^21,23,31,36^. In pioneering experiments, Debarbieux et al. investigated the impact of phage dose relative to bacterial burden, i.e. the multiplicity of infection (MOI), in a murine pneumonia model^37^. Mice treated at a phage:bacteria ratio of 1:10 died within 5 days, whereas >80% survival was achieved at a ratio of 1:1 and 100% survival at 10:1. The importance of phage:bacteria ratios was further supported by Lin et al. in a murine bacteremia model^18^. In that study, mice infected with 5 × 10^8^ CFU of multidrug-resistant *Pseudomonas aeruginosa* achieved rapid bacterial clearance once phage doses reached at least 10^5^ PFU/mouse, whereas higher doses did not further improve efficacy. The minimal phage concentration required to reduce the bacterial population might therefore define a productive infection threshold for active phage therapy. Importantly, both insufficient and excessive phage concentrations may be suboptimal: low titers may fail to initiate productive infection, whereas excessively high titers could impair productive infection^21,38^.

If MOI was a key determinant of efficacy, then achieving a sufficient C_max_ at the infection site may be critical to exceed this productive infection threshold. Conversely, if sustained phage exposure is required, cumulative exposure over time (AUC) may represent a more relevant PK/PD surrogate. It remains unclear whether lysis from without contributes to passive phage therapy at very high titers, where extreme MOIs may exceed the abortive infection threshold^39,40^.

In this context, accurate pharmacokinetic modeling becomes essential. The second-order pharmacokinetic behavior observed here has two major consequences: in comparison with first-order estimates, it reduces the expected exposure over time (AUC) even if high peak concentrations (C_max_) were to be achieved and limits systemic accumulation after repeated dosing. Accordingly, repeated administration increased overall exposure, although to a much lesser extent than would have been expected under first-order elimination. Local retention may permit transient buildup in peripheral compartments, but the present data suggest that this would require very short dosing intervals and high titers. Moreover, any such effect might be offset if clearance accelerates following activation of the immune system^20^. These effects might critically influence therapeutic efficacy, particularly if phage activity depends on cumulative exposure.

Distribution analyses further revealed that the route of administration is another major determinant of therapeutic phage exposure. Intravenous delivery resulted in rapid systemic dissemination but relatively limited peak titers in the peripheral (tissue cage) compartment. In contrast, direct local administration achieved substantially higher concentrations at the target site. Combined intravenous and local administration is also frequently used empirically^10^, yet our data showed no relevant additive pharmacokinetic benefit compared with local administration alone. Intraperitoneal delivery produced comparatively low systemic exposure, whereas subcutaneous administration failed to generate detectable titers. Together, these findings indicate that anatomical access, rather than dose alone, governs effective phage delivery to sites of infection. Accordingly, achieving a therapeutically relevant inundation threshold depends primarily on local peak concentrations, supporting targeted delivery strategies for compartmentalized infections.

The present phage panel encompassed major tailed-phage morphotypes and included representatives of two widely used therapeutic phage genera, all of which consistently exhibited second-order elimination kinetics and differed primarily in their rate constants. This conserved behavior supports the generalizability of our findings within rats and suggests that unified models may predict phage pharmacokinetics across different phage types and therapeutic contexts. Validation using published data further supports generalizability of these findings across additional phages, doses, and animal species. Extrapolation from animals to humans, however, requires caution, and highlights the need for phase I trials in humans integrating intensive pharmacokinetic, immunological, and microbiological measurements. Such studies will be essential to validate these findings, characterize inter-individual variability, and define exposure–response relationships. These data are increasingly required within regulatory frameworks for biologics to support rational dose selection and clinical development. Integrating pharmacokinetic modeling into early clinical studies will therefore be critical to transition phage therapy from empirical use to a quantitatively guided, mechanism-based treatment strategy. Further translational research should also identify phage-binding receptors on the cell types responsible for clearing circulating bacteriophages and characterize the kinetics of phage binding, internalization, and degradation. This may clarify whether saturation of the elimination system is potentially achievable and could reveal possible targets to improve phage exposure at sites of infection.

In summary, phage pharmacokinetics are governed by second-order, concentration-dependent clearance, while administration route and dosing frequency shape systemic and peripheral exposure. The conservation of these dynamics across diverse phages supports unified models to predict phage behavior across therapeutic contexts. More broadly, variability in phage exposure appears to be driven primarily by host factors rather than phage identity. Together, these findings provide a foundation for rational dose design and optimization of phage targeting bacterial pathogens.

## METHODS

### Key resource table

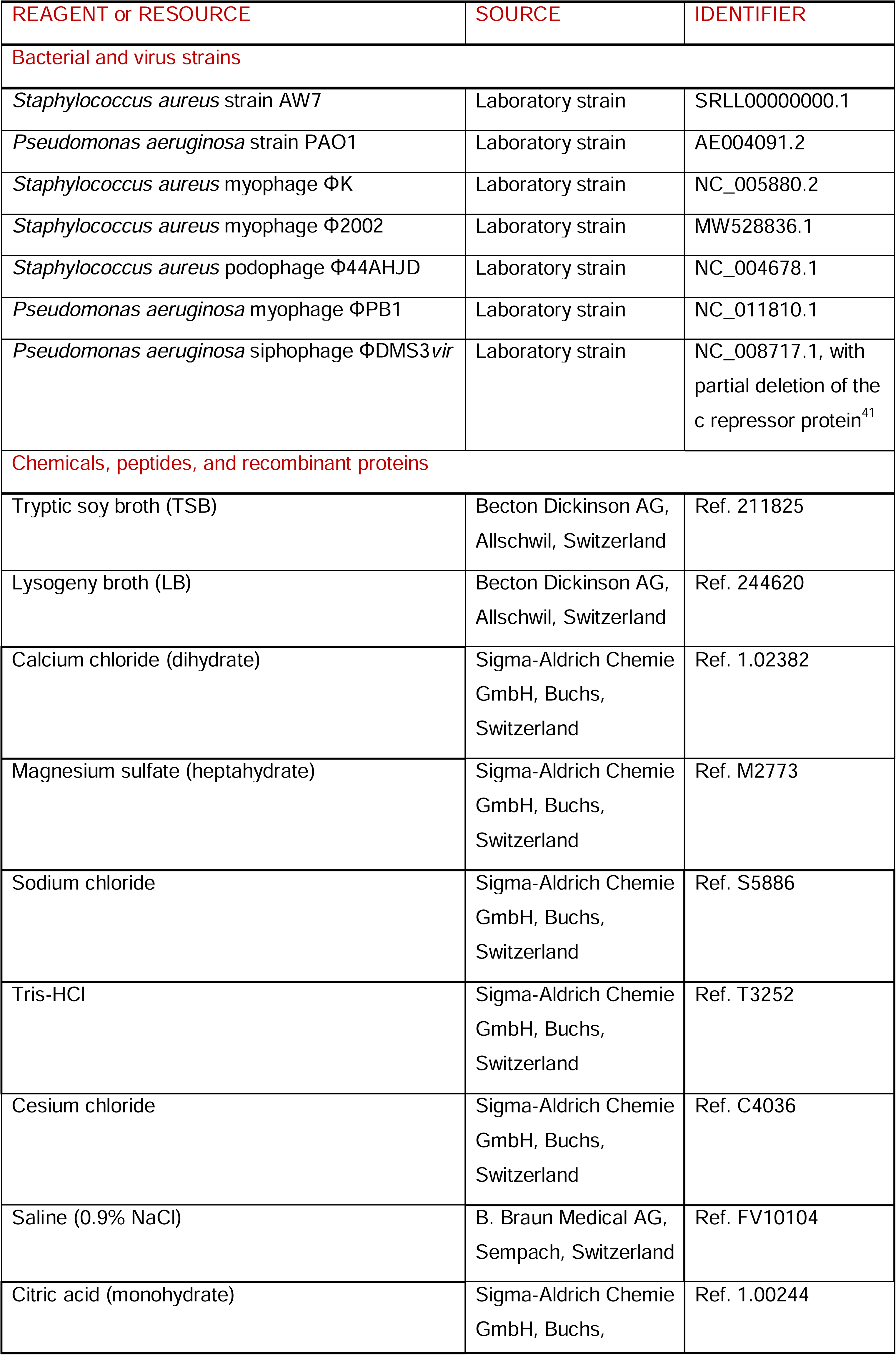
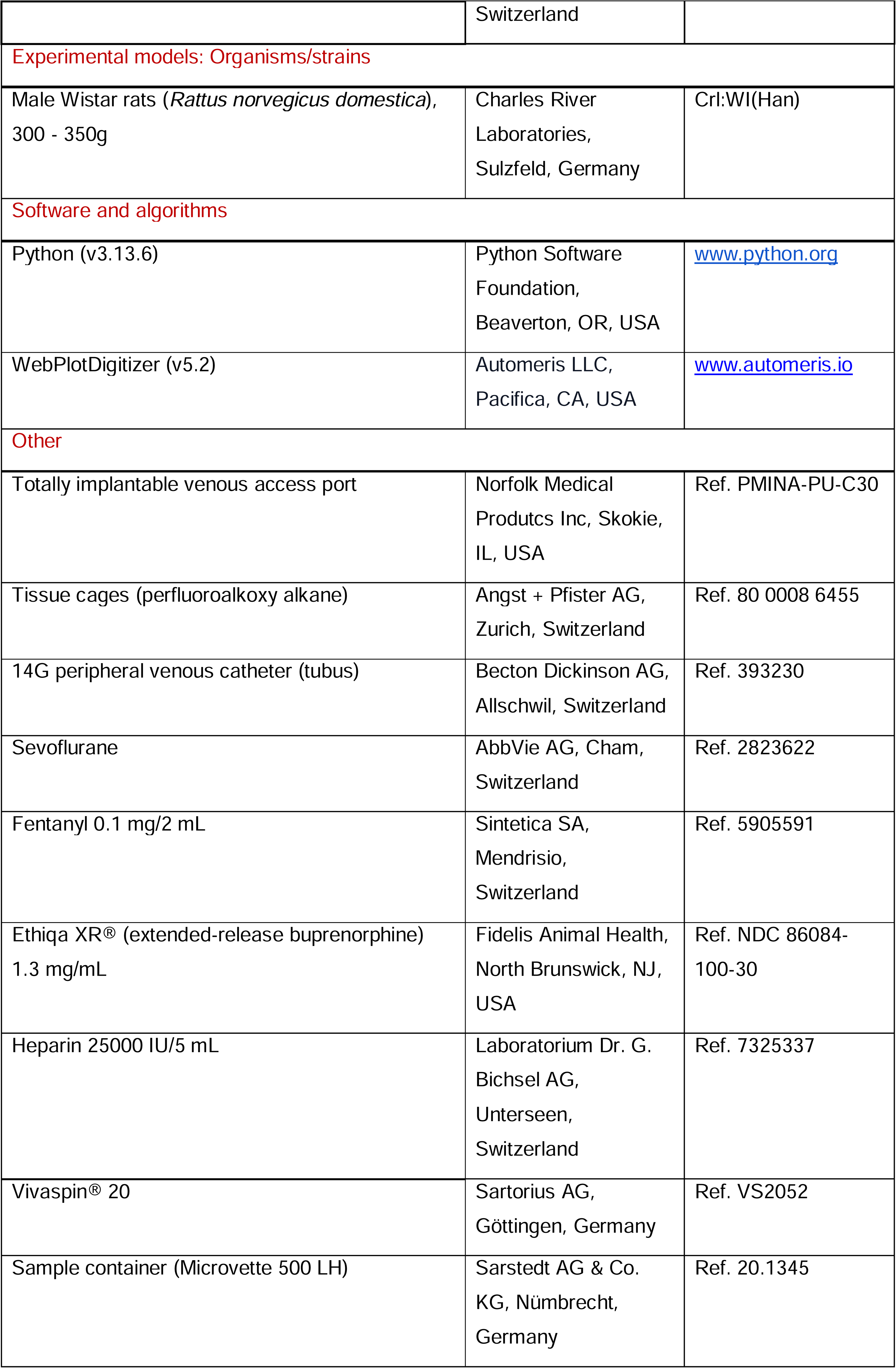

### Animal model and ethical approval

Male Wistar rats (Crl:WI(Han), 300-350g; Charles River Laboratories) were housed under specific pathogen-free conditions with a 12-h light/dark cycle at 23°C ± 1°C, with food and water provided ad libitum. All procedures were conducted in accordance with the Swiss Animal Protection Law and approved by the Cantonal Committee on Animal Experiments of the State of Bern (Authorizations BE139/2022, CH35499). Humane endpoints included respiratory distress, sopor or seizures, loss of reflexes or inability to remain upright, and >10% body weight loss.

### Surgical procedures

Following induction with sevoflurane and intraperitoneal injection of 20 μg/kg fentanyl, animals were intubated with a 14 G peripheral venous catheter (Becton Dickinson AG). Under balanced intubation anesthesia (sevoflurane and fentanyl), animals were then equipped with a total implantable venous access port (TIVAP) via the right external jugular vein and the tip at the cavoatrial junction. In addition, two perforated perfluoroalkoxy alkane tissue cages (outer measures 32 × 10 mm, inner measures 8.5 x 27.5 mm; Angst + Pfister AG) were implanted subcutaneously on the dorsum (Figure 1A). Postoperative analgesia with extended-release buprenorphine (Ethiqa XR®) was administered according to body weight. Animals were monitored daily using a score sheet and allowed to recover from surgery for three weeks before pharmacokinetic studies, enabling implant compartmentalization through foreign-body reaction with encapsulation. Meanwhile, TIVAP were locked with heparin-citrate (500 IU/mL, 4% in 0.9% NaCl) and changed at minimum weekly or after each use, respectively (Figure 1C).

### Study design

Pharmacokinetic experiments were designed to evaluate (i) systemic elimination following single-dose intravenous (IV) administration; (ii) distribution between the central (blood) and peripheral (tissue cage fluid, TCF) compartments following IV, local (intra-tissue cage, TC), intraperitoneal (IP), or subcutaneous (SC) administration; and (iii) accumulation during repeated dosing via IV, local, or combined routes. Primary outcome measures were phage titers in blood and TCF. Secondary outcome measures were peak concentrations in compartments (C_max_) and exposure over 48h expressed as area under the curve (AUC_48h_). Doses of 5 × 10^9^ PFU, corresponding to 0.25 mL of a phage preparation with a titer of 2 × 10^10^ PFU/mL, were administered, except for local administration into tissue cages (5 × 10^7^ PFU, or 0.25 mL of a phage preparation with a titer of 2 × 10^8^ PFU/mL). For repeat dosing experiments, doses were administered at 0 h, 4 h, 8 h, and 24 h. Catheters were flushed with 0.5 mL of 0.9% NaCl after each administration. Samplings occurred at baseline (before injection) and after 5 min, 2 h, 4 h, 8 h, 24 h, and 48 h. Microvettes 500 LH (Sarstedt AG & Co. KG) were used as a sample container. All injections and samplings were performed under brief sedation with sevoflurane.

### Bacterial culture

*S. aureus* strain AW7 and *P. aeruginosa* strain PAO1 were revived from 20% (v/v) glycerol stocks and streaked onto tryptic soy agar (TSA) and lysogeny broth (LB) agar plates, respectively. Plates were incubated at 37°C for 18 hours and stored at 4°C for a maximum of two weeks. For experiments, single colonies were used to inoculate 10 mL of the corresponding liquid medium (TSB or LB), followed by overnight incubation at 37°C with orbital shaking at 200 rpm.

### Phage preparation

The five virulent phages were propagated on their respective host strains (*S. aureus* AW7, *P. aeruginosa* PAO1, Key resource Table) in semi-solid agar overlays after cross-contamination was excluded by sequencing. Briefly, overnight bacterial cultures were mixed with phages (multiplicity of infection 0.1) and incorporated into soft agar (0.38%) prepared in TSB (*S. aureus*) or LB (*P. aeruginosa*) supplemented with CaCl_2_ (2 mM) and, for pseudomonal phages, MgSO_4_ (10 mM). After incubation at 37°C for 18 h, the overlay was harvested, resuspended in liquid medium, clarified by centrifugation (10,000 × g, 15 min), and filtered (0.22 µm) to obtain crude lysates. Phages were purified by cesium chloride (CsCl) density-gradient ultracentrifugation (1.31–1.57 g cm^-3^; 100 000 × g, 2 h, 4°C). Phage bands were collected and further purified using 300 kDa ultrafiltration devices (Vivaspin® 20) through three sequential dilution-concentration cycles (20 mL to 200 µL) into SM buffer. Phage suspensions were normalized to 2 × 10^10^ PFU/mL and stored at 4°C for a maximum of two weeks.

### Phage quantification

Phage titers were determined using the double-layer agar assay^42^. To allow quantification of high titers, 10 µL of each sample was serially diluted (10^-1^ to 10^-8^) and spotted in technical triplicate (3 × 4 µL per dilution) onto TSB (*S. aureus*) or LB (*P. aeruginosa*) plates overlaid with soft agar (0.75%) containing the appropriate host strain at a final concentration of 10^8^ CFU/mL and supplemented with CaCl_2_ (2 mM) and, for pseudomonal phages, additionally with MgSO_4_ (10 mM). To improve sensitivity for low titers, an additional assay was performed in parallel, in which 50 µL of undiluted sample was incorporated directly into the overlay, resulting in a detection limit of 20 PFU/mL. Plates were incubated at 37°C for 18 hours before enumeration of plaque-forming units (PFU).

### Pharmacokinetic analysis

Phage titers were log_10_-transformed for analysis. Zero-, first-, and second-order kinetic models were compared using observed median data and interquartile ranges (IQR). Simulations and estimations were based on the appended formulas (Supplemental Table 1). Half-life for first-order kinetics was estimated based on a corresponding decay from mean peak to mean trough titers across all phages. For second-order kinetics, rate constants were extracted from the slope of the linear regression between inverse concentration and time and model fit was assessed using the coefficient of determination (*R^2^*). *n* represents biological replicates.

### Pharmacokinetic parameters

Maximum concentration (C_max_) and area under the concentration–time curve over 48 h (AUC_48h_) were derived from median titers. Calculation of observed and predicted AUC_48h_ was based on the trapezoidal rule. Values below the limit of detection were treated as 20 PFU/mL.

### Extraction of published data

Relevant studies reporting mammalian in vivo pharmacokinetics following intravenous bacteriophage administration were identified through searches of MEDLINE using generic terms and the reference lists of relevant articles. Eligible datasets were required to provide systemic phage titer data over time. Datasets were excluded if more than 50% of values were at the limit of detection (LOD) or if the experimental conditions were not considered comparable, for example in irradiated animals^32^. Systemic phage titer data were extracted from published figures using WebPlotDigitizer (v5.2). Values reported at or below the LOD were treated as missing to avoid biasing model fitting.

### Statistical analysis

Models were fitted separately to the data, with each curve anchored to the first included timepoint. Model trajectories were plotted through the last included timepoint. For second-order kinetics, rate constants were derived from the slope of inverse concentration versus time. The performance of zero-, first-, and second-order models was compared using the root mean square error (RMSE) of log_10_-transformed titers to quantify the average prediction error. Relative model fit was assessed using the corrected Akaike information criterion (AICc) with k = 1, calculated from residuals between model predictions and the median observed log_10_-transformed titers, to identify the best-predicting model. For each dataset, ΔAICc was calculated by subtracting the lowest AICc among the models, with ΔAICc = 0 indicating the best-fitting model. Differences in ΔAICc across zero-, first-, and second-order models were assessed using the Friedman test. Pairwise comparisons between models were performed using two-sided Wilcoxon signed-rank tests with Holm correction for multiple testing. All analyses were performed using Python (v3.13.6).

## RESOURCE AVAILABILITY

### Lead contact

Requests for further information and resources should be directed to and will be fulfilled by the lead contact, Yok-Ai Que.

## Materials availability

This study did not generate any new, unique reagents.

## Data and code availability

Data, code and any additional information required to reanalyze the data reported in this paper is available from the lead contact upon request.

## ACKNOWLEDGEMENTS

The authors thank Sandra Nansoz and Tessa Guggiari for their excellent technical assistance.

## AUTHORS CONTRIBUTIONS

J.L.G., D.R.C. and Y.-A.Q. conceived and designed the study. J.L.G, B.G., J.P. and C.B performed the experiments. J.L.G. and D.R.C. analyzed the data. J.L.G., J.P. and Y.-A.Q. wrote the manuscript. All authors reviewed and revised the manuscript for intellectual content and approved the final version.

## SUPPLEMENTAL INFORMATION

Supplemental Figures S1–S3 and Supplemental Tables S1 and S2

## DECLARATION OF GENERATIVE AI AND AI-ASSISTED TECHNOLOGIES IN THE WRITING PROCESS

During the preparation of this work the authors used ChatGPT in order to improve the readability and language of the manuscript. The authors reviewed and edited the output as needed and take full responsibility for the content of the published article.

---

**TABLE 1:** Observed and model-predicted systemic phage exposure (AUC_48h_). Observed and predicted areas under the concentration–time curve over 48 h (AUC_48h_). Predicted AUC_48h_ were calculated from the corresponding zero-, first-, and second-order models of Figure 3. Values are plaque forming units (PFU)×h/mL.

| Phage |  | Observed AUC<sub>48h</sub> | Predicted AUC<sub>48h</sub> for |  |  |
| --- | --- | --- | --- | --- | --- |
|  |  |  | Zero-order | First-order | Second-order |
| *S. aureus* | **ΦK** (*n* = 9) | $9.11 \times 10^6$ | $2.22 \times 10^8$ | $5.51 \times 10^7$ | $9.01 \times 10^+$ |
| | **Φ2002** (*n* = 9) | $4.89 \times 10^4$ | $1.09 \times 10^6$ | $2.70 \times 10^5$ | $4.85 \times 10^4$ |
| | **Φ44AHJD** (*n* = 3) | $2.52 \times 10^5$ | $4.01 \times 10^6$ | $9.93 \times 10^5$ | $3.48 \times 10^5$ |
| *P. aeruginosa* | **ΦPB1** (*n* = 8) | $2.55 \times 10^7$ | $6.40 \times 10^8$ | $1.56 \times 10^8$ | $2.56 \times 10^7$ |
| | **ΦDMS3vir** (*n* = 6) | $4.77 \times 10^7$ | $1.25 \times 10^9$ | $2.94 \times 10^8$ | $4.78 \times 10^7$ |

