## Supplemental for "Second-order kinetics describe systemic clearance of therapeutic bacteriophages"

Prof. Dr. Yok-Ai Que, MD, PhD

Department of Intensive Care Medicine, INO E-419

Inselspital; Bern University Hospital, University of Bern

CH-3010 Bern, Switzerland

**INDEX OF SUPPLEMENTAL MATERIAL**

| **Supplemental Table 1** | Theoretical rate laws and kinetic relationships for zero-, first-, and second-order elimination | p. 2 |
| --- | --- | --- |
| **Supplemental Figure 1** | Theoretical elimination kinetics and calculation of rate constants | p. 3 |
| **Supplemental Figure 2** | Comparison of published phage blood kinetic data | p. 4 |
| **Supplemental Table 2** | Overview of the experiments included in the comparison of published phage blood kinetic data | p. 6 |
| **Supplemental Figure 3** | Route-dependent phage exposure in blood following single-dose administration of ΦK | p. 7 |
| **References** |  | p. 8 |

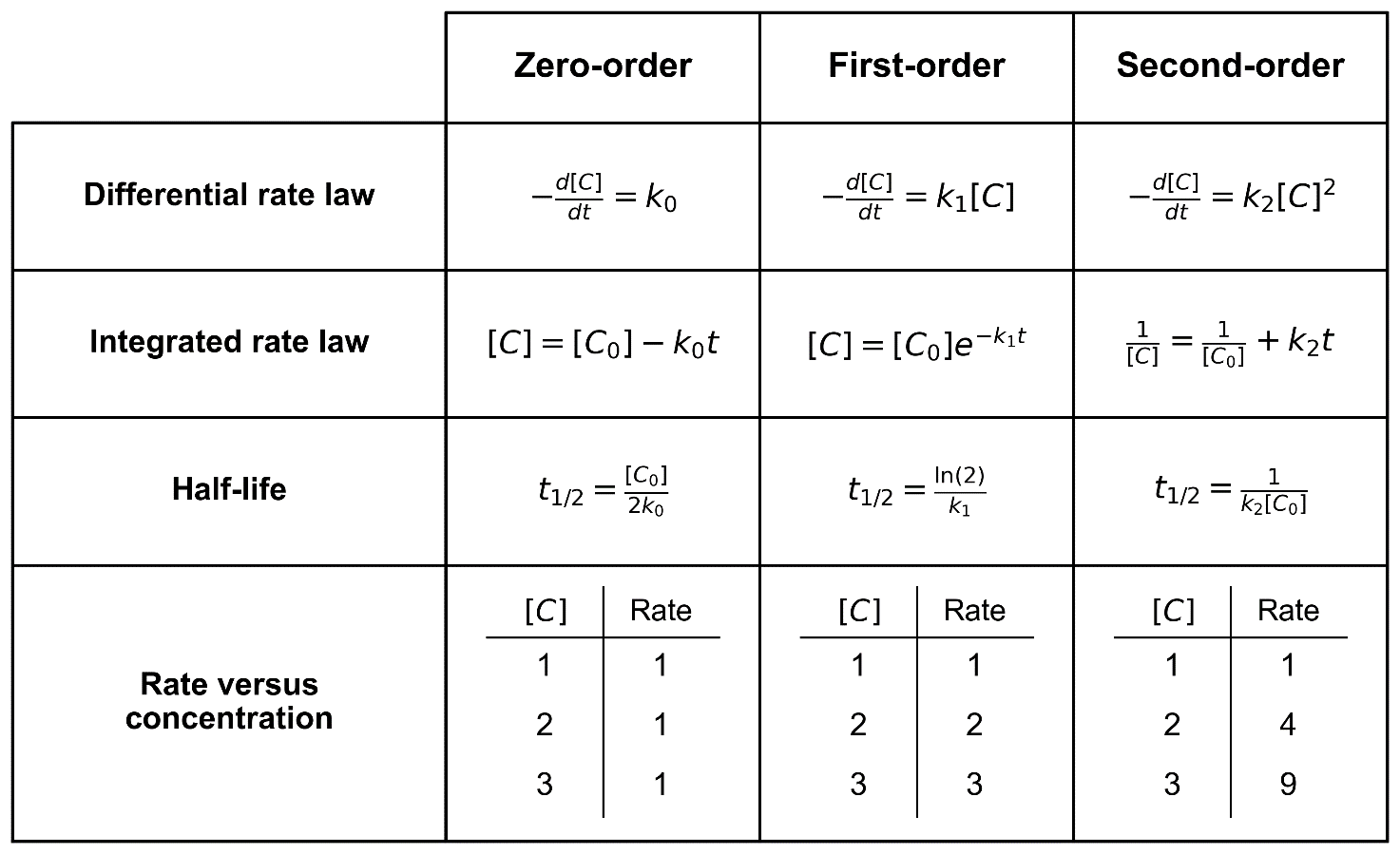

**SUPPLEMENTAL TABLE 1: Theoretical rate laws and kinetic relationships for zero-, first-, and second-order elimination.**

This table summarizes the fundamental equations describing zero-order, first-order, and second-order kinetics. For each kinetic order, the differential rate law, integrated rate law, and corresponding half-life expression are shown. Elimination kinetics differ in how the rate depends on concentration: zero-order rates are concentration independent, first-order rates increase linearly with concentration, and second-order rates increase with the square of concentration. The table also illustrates how reaction rate varies with concentration for each kinetic order. Together with Supplemental Figure 1, these relationships provide the theoretical framework for identifying the order of elimination.

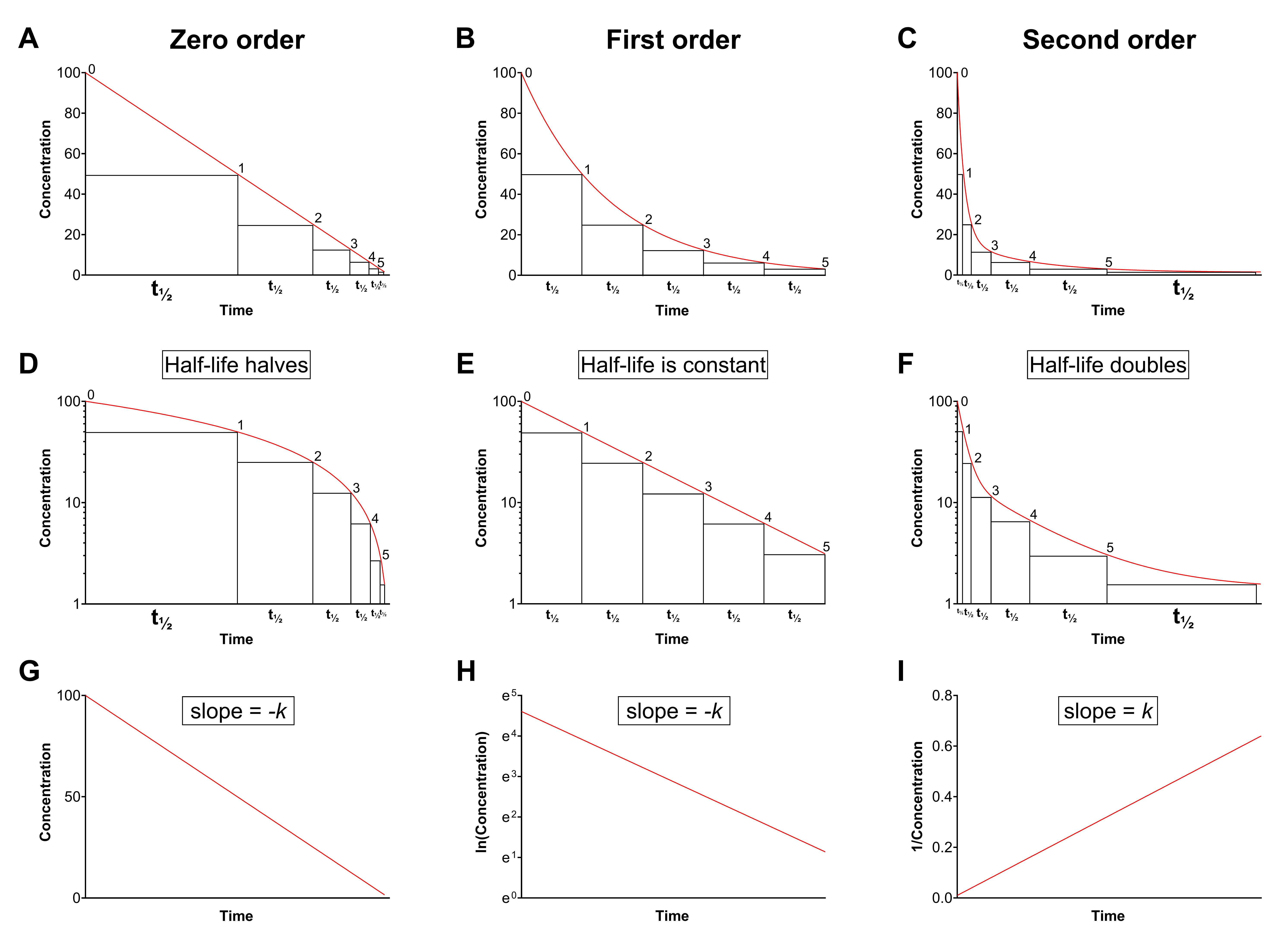

**SUPPLEMENTAL FIGURE 1: Theoretical elimination kinetics and calculation of rate constants.**

The figure illustrates zero-order (left column), first-order (middle column), and second-order (right column) elimination kinetics. (**A–C**) Concentration–time curves on linear axes show how the half-life changes with each kinetic order. In zero-order kinetics the half-life decreases progressively (halves), in first-order kinetics the half-life remains constant, and in second-order kinetics the half-life increases progressively (doubles). (**D–F**) The same relationships are shown on semi-logarithmic axes. Importantly, first-order kinetics linearize on a logarithmic concentration scale, whereas second-order kinetics do not. (**G–I**) Plots used to determine the elimination rate constant (*k*). For zero-order kinetics, concentration plotted against time yields a straight line with slope −*k_0_*. For first-order kinetics, ln(concentration) plotted against time gives a straight line with slope to −*k_1_*. For second-order kinetics, inverse concentration plotted against time results in a straight line with slope *k_2_*. Kinetic order can be identified from the plot that most closely approximates linearity, and the corresponding rate constant is calculated from the slope of the linear regression.

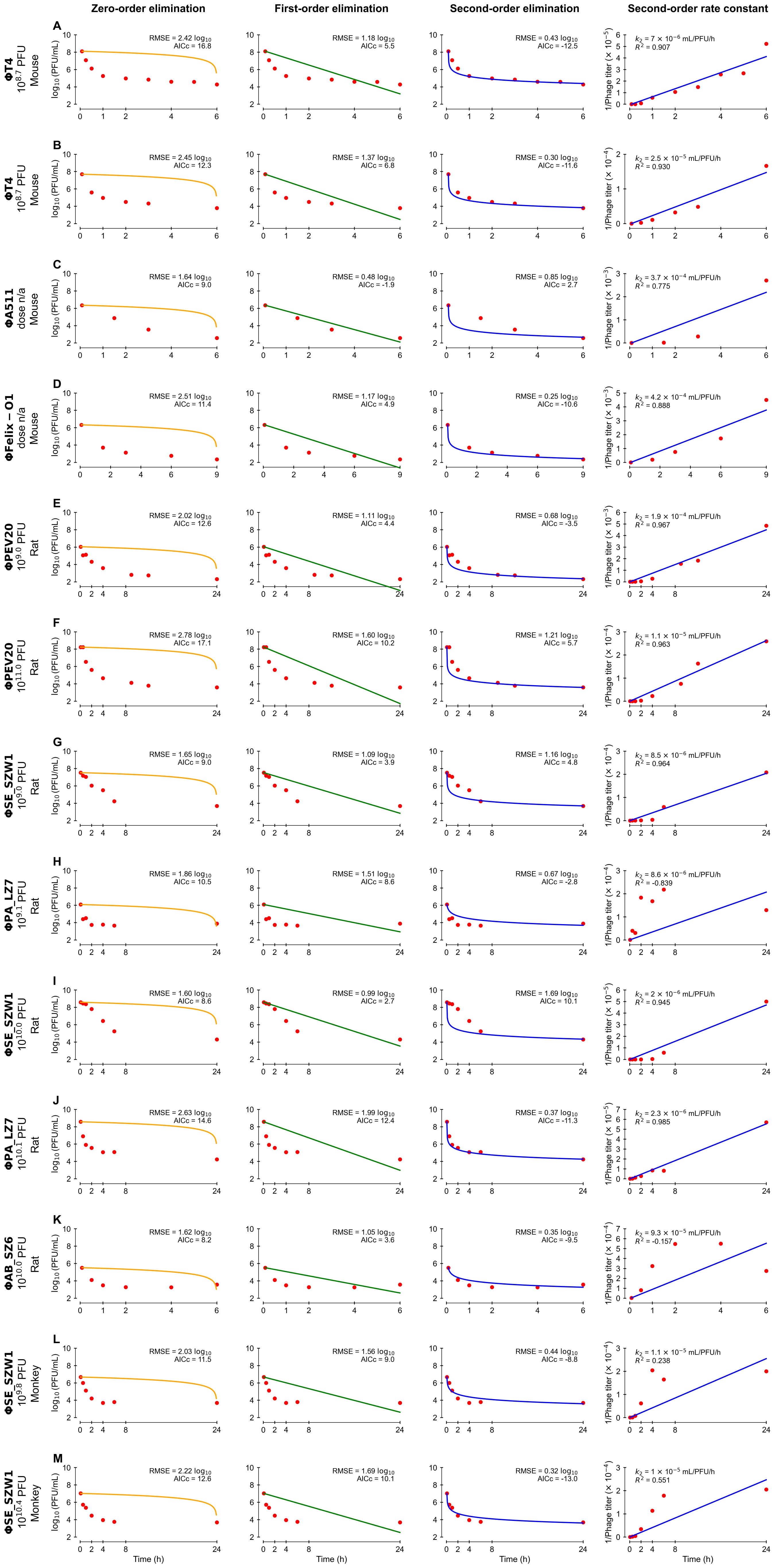

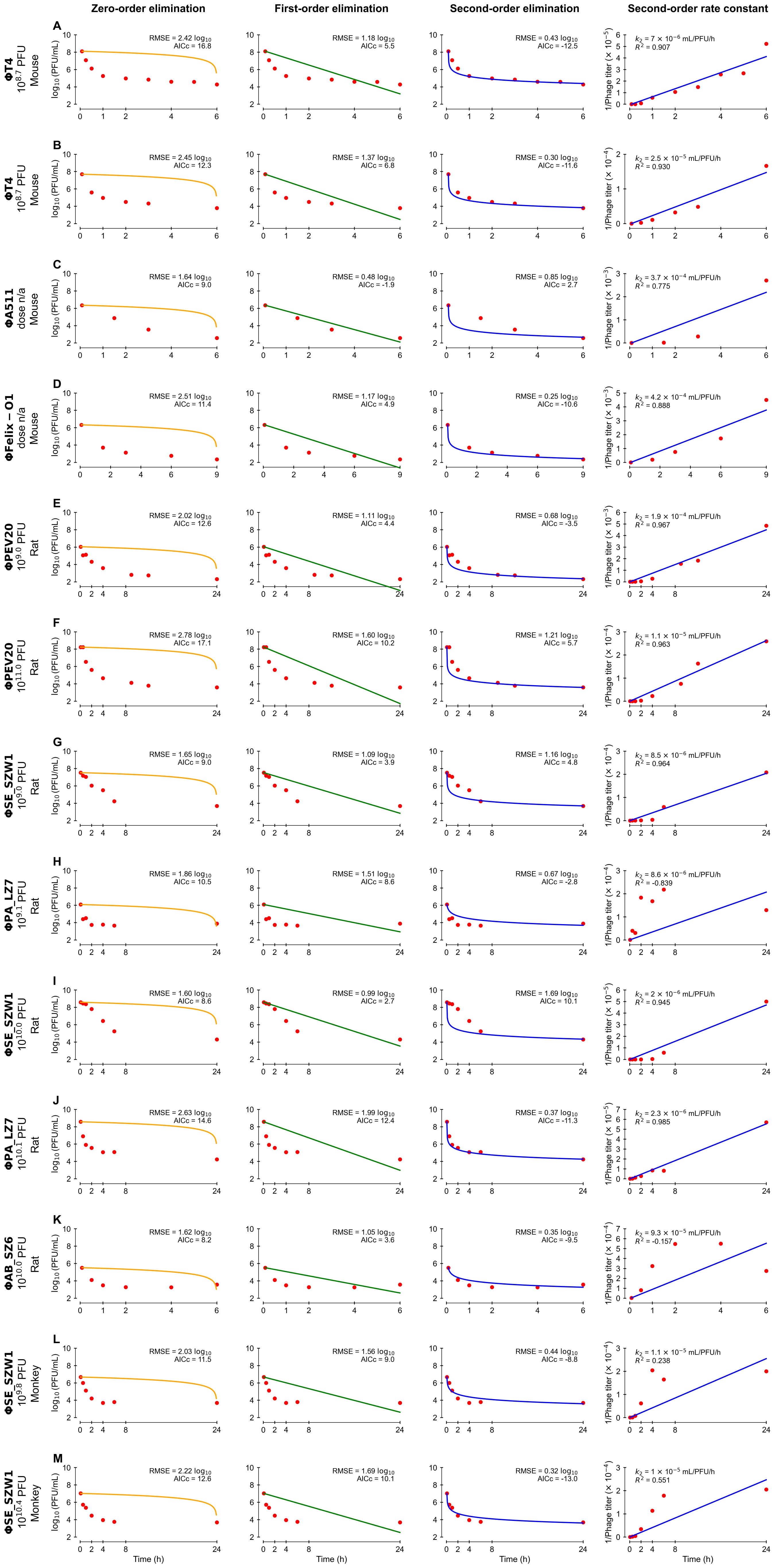

**SUPPLEMENTAL FIGURE 2: Comparison of published phage blood kinetic data.**

Model fits (colored lines) to observed median titers (red datapoints) extracted from published experiments (Supplemental Table 2). For each experiment, zero-order (orange), first-order (green), and second-order (blue) fits are shown. The corresponding linear regressions of inverse phage titer versus time are displayed in blue, with *k_2_* specifying the second-order rate constant (slope) and *R^2^* showing regression fit. The root mean square error (RMSE) and corrected Akaike information criterion (AICc), calculated from log_10_-transformed titers, indicate the average prediction error and relative model fit, respectively.

In most experiments, second-order elimination provided a good fit and outperformed first-order elimination (Figure 4). However, this re-analysis is limited by the quality and heterogeneity of the published source data. Limits of detection, animal numbers, body weight, and sex were not consistently reported. Handling of values below the LOD varied markedly between studies. Most datasets reported means, which are sensitive to skewness and outliers, particularly in small groups.

| **Study** | **Animal** | **n** | **Sex** | **Phage** | **Morphotype** | **Bacterial host** | **Dose (PFU)** | **Titer measure** | **LOD (PFU/mL)** |
| --- | --- | --- | --- | --- | --- | --- | --- | --- | --- |
| Inchley 1969^1^ | Mouse | 3 | N/A | ΦT4 | Myophage | *Escherichia coli* | 5 × 10^8^ | Mean | N/A |
| Inchley 1969^1^ | Mouse | 4 | N/A | ΦT4 | Myophage | *Escherichia coli* | 5 × 10^8^ | Mean | N/A |
| Kim 2008^2^ | Mouse | 8 | Male | ΦA511 | Myophage | *Listeria monocytogenes* | N/A | Mean | 200 |
| Kim 2008^2^ | Mouse | 8 | Male | ΦFelix-O1 | Myophage | *Salmonella enterica* | N/A | Mean | 200 |
| Lin 2020^3^ | Rat | 3 | Male | ΦPEV20 | Podophage | *Pseudomonas aeruginosa* | 3 × 10^9^ | Geometric mean | 50 |
| Lin 2020^3^ | Rat | 3 | Male | ΦPEV20 | Podophage | *Pseudomonas aeruginosa* | 3 × 10^11^ | Geometric mean | 50 |
| Tan 2024^4^ | Rat | 5 | Female | ΦSE_SZW1 | Siphophage | *Salmonella enterica* | 5 × 10^9^ | Mean | 10^3^ |
| Tan 2024^4^ | Rat | 6 | Mixed | ΦPA_LZ7 | Myophage | *Pseudomonas aeruginosa* | 5 × 10^9^ | Mean | 10^3^ |
| Tan 2024^4^ | Rat | 5 | Female | ΦSE_SZW1 | Siphophage | *Salmonella enterica* | 5 × 10^10^ | Mean | 10^3^ |
| Tan 2024^4^ | Rat | 5 | Mixed | ΦPA_LZ7 | Myophage | *Pseudomonas aeruginosa* | 5 × 10^10^ | Mean | 10^3^ |
| Tan 2024^4^ | Rat | 5 | Female | ΦAB_SZ6 | Podophage | *Acinetobacter baumanii* | 5 × 10^10^ | Mean | 10^3^ |
| Tan 2024^4^ | Macaque | 4 | Mixed | ΦSE_SZW1 | Siphophage | *Salmonella enterica* | 6 × 10^9^ | Mean | 10^3^ |
| Tan 2024^4^ | Macaque | 6 | Mixed | ΦSE_SZW1 | Siphophage | *Salmonella enterica* | 2 × 10^10^ | Mean | 10^3^ |

**SUPPLEMENTAL TABLE 2: Overview of the experiments included in the comparison of published phage blood kinetic data.**

Summary of the 13 experiments extracted from four studies that were used for comparison of published phage blood-kinetic data (Supplemental Figure 2). Presented variables include animal and phage characteristics, dose, and reported measure of central tendency (titer measure). Datasets were excluded when more than half of the values were below the LOD or when the experimental condition was potentially not comparable, such as irradiated mice^1^. LOD indicates the limit of detection, N/A stands for not available, and PFU for plaque-forming units.

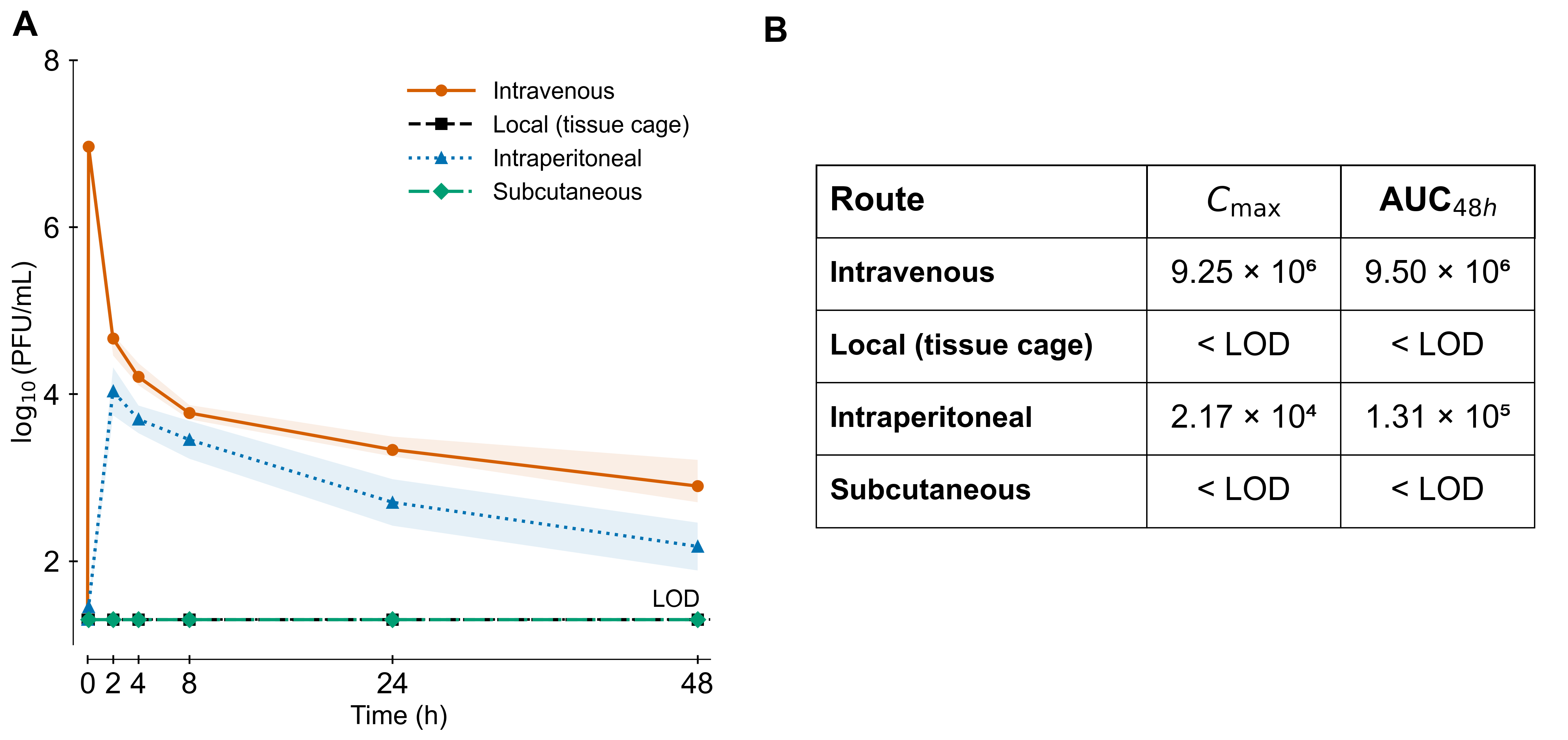

**SUPPLEMENTAL FIGURE 3: Route-dependent phage exposure in blood following single-dose administration of ΦK.**

(**A**) Phage titers in blood following administration of ΦK via different routes: intravenous (n = 18), local (tissue cage; n = 3), intraperitoneal (n = 4), and subcutaneous (n = 2). Doses were 5 × 10^9^ plaque forming units (PFU) for intravenous administration and 5 × 10^7^ PFU for local tissue cage administration. Data points represent median titers, and shaded areas indicate the interquartile range (IQR). The dotted horizontal line indicates the limit of detection (LOD; 20 PFU/mL). (**B**) Maximum concentration (C_max_), and systemic exposure (AUC_48h_) in blood calculated from median titers over 48 h using the trapezoidal rule.
